# Neutralization sensitivity of the SARS-CoV-2 Omicron BA.2.75 sublineage

**DOI:** 10.1101/2022.08.04.502609

**Authors:** Henning Gruell, Kanika Vanshylla, Pinkus Tober-Lau, David Hillus, Leif Erik Sander, Florian Kurth, Florian Klein

## Abstract

The recently emerged BA.2.75 Omicron sublineage of SARS-CoV-2 identified in numerous countries is rapidly increasing in prevalence in regions of India. Compared with BA.2, the spike protein of BA.2.75 differs in nine amino acid residues. To determine the impact of the spike mutations on polyclonal and monoclonal antibody activity, we investigated the neutralization sensitivity of BA.2.75 in comparison with B.1, BA.2, BA.2.12.1, and BA.4/5. Analysis of post-boost samples from 30 vaccinated individuals revealed significantly lower serum neutralizing activity against BA.2.75 than against BA.2. However, BA.2.75 was more sensitive to serum neutralization than the widely circulating BA.4/5 sublineages. Moreover, evaluation of 17 clinical-stage monoclonal antibodies demonstrated individual differences in Omicron sublineage activity. Notably, some authorized antibodies with low activity against other Omicron sublineages demonstrated high BA.2.75 neutralizing potency. Our results indicate a less pronounced degree of antibody evasion of BA.2.75 compared with BA.4/5 and suggest that factors beyond immune evasion may be required for an expansion of BA.2.75 over BA.4/5.

## Text

On July 15, 2022, the European Centre for Disease Prevention and Control (ECDC) elevated the SARS-CoV-2 Omicron BA.2.75 sublineage to the category of “variant of interest”. After its first identification in a sample collected at the end of May 2022, genomic surveillance revealed a rapid increase in the proportion of BA.2.75 amongst sequenced SARS-CoV-2 infections with up to >20% in regions of India by mid-July 2022.^1^ Moreover, cases of BA.2.75 infections have been reported in numerous countries across the globe.^1^ Compared with the parental BA.2 lineage of SARS-CoV-2, the spike protein of BA.2.75 differs in nine amino acid residues in the N-terminal domain (K147E, W152R, F157L, I210V, G257S) and the receptor binding domain (D339H, G446S, N460K, R493Q) (**Figure S1**). By affecting critical epitopes, mutations in these domains can confer a growth advantage through reduced susceptibility to SARS-CoV-2 neutralizing antibodies.^2^ To investigate antibody sensitivity of BA.2.75 in comparison to prevalent Omicron sublineages, we performed neutralization assays using pseudoviruses expressing the B.1 (D614G), BA.2, BA.4/5, BA.2.12.1, or BA.2.75 spike proteins.

First, we determined fifty-percent serum inhibitory dilutions (ID_50_s) in serum samples collected 4 weeks after a BNT16b2 vaccine booster dose in a cohort of 30 health care workers and elderly individuals (**Table S1**). All individuals had previously received two BNT162b2 doses and no intermittent SARS-CoV-2 infections were reported. While neutralizing activity against Omicron sublineages was considerably lower than against B.1, differences between individual Omicron sublineages were more subtle (geometric mean ID_50_s of 6,429 against B.1, 1,725 against BA.2, 615 against BA.4/5, 1,052 against BA.2.12.1, and 1,203 against BA.2.75) (**Figure 1A** and **Figure S2**). Compared with the other Omicron sublineages, serum activity against BA.2.75 was significantly lower than against BA.2 (*p*=0.0145) but higher than against BA.4/5 (*p*=0.0329) (**Figure S2**).

**Figure 1.**
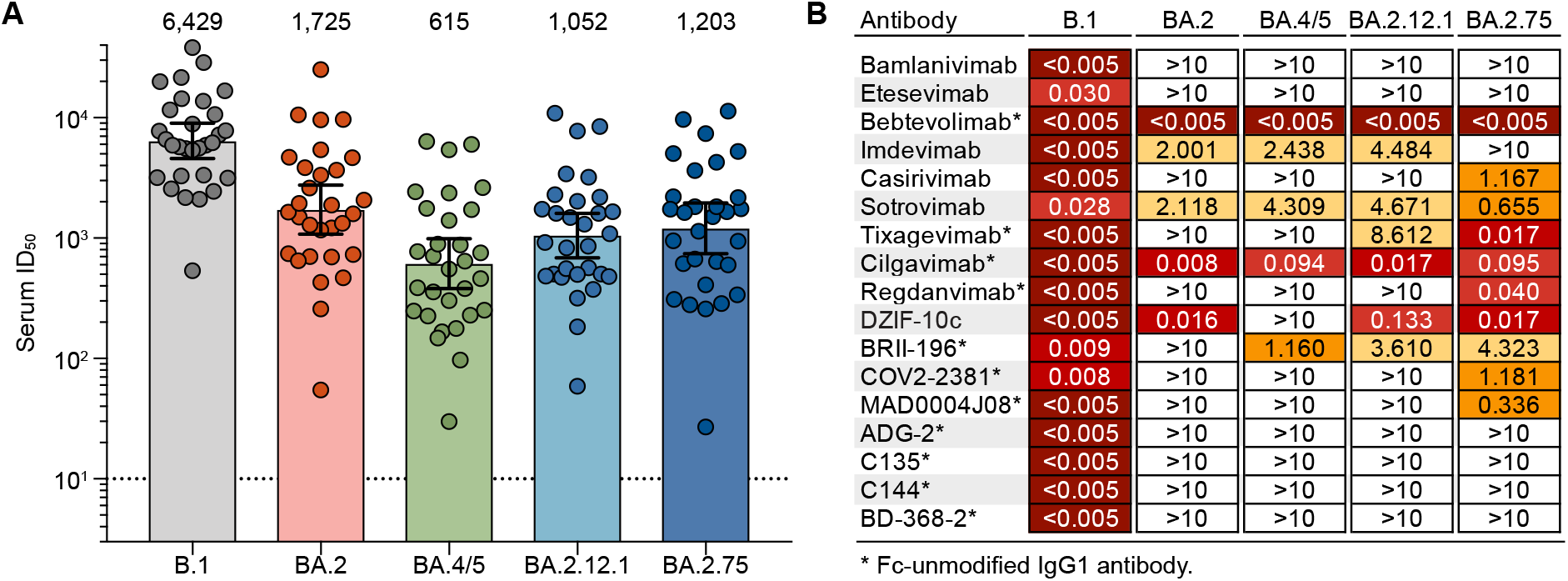
Neutralization sensitivity of the BA.2.75 sublineage. **(A)** Fifty-percent inhibitory dilutions (ID50s) against SARS-CoV-2 variants in samples collected four weeks after a BNT162b2 booster vaccination (*n*=30) determined by pseudovirus neutralization assay. Circles indicate average ID50s of two experiments for each variant and individual participant. Bars and numbers indicate geometric mean ID50s. Solid lines indicate 95% confidence intervals and the dashed line shows the lower limit of quantification. **(B)** Fifty-percent inhibitory concentrations (IC50s) of monoclonal antibodies against SARS-CoV-2 variants determined by pseudovirus neutralization assay (average IC50 of two experiments for each variant).

Next, we investigated the activity of 17 monoclonal antibodies conditionally authorized for the use against COVID-19 or in advanced stages of clinical investigation by determining their fifty-percent inhibitory concentrations (IC_50_s). Most antibodies failed to neutralize either BA.2, BA.4/5, or BA.2.12.1 (IC_50_ >10 µg/ml). Notably, however, several of these antibodies demonstrated appreciable activity against BA.2.75 (**Figure 1B** and **Figure S3**). For example, tixagevimab and regdanvimab, authorized for COVID-19 prevention and treatment (in South Korea), respectively, showed no or only poor activity against any of the other tested Omicron sublineages but potently neutralized BA.2.75 (IC_50_ ≤0.04 µg/ml). As against all other Omicron sublineages, the antibody bebtelovimab also demonstrated potent BA.2.75 neutralizing activity. In total, while only 30-35% of the tested antibodies neutralized BA.2, BA.4/5, or BA.2.12.1, activity against BA.2.75 was detected for 59% of antibodies.

Variants of SARS-CoV-2 with reduced sensitivity to neutralizing antibodies can pose a challenge to immunity induced by vaccination and/or infection and render therapeutic monoclonal antibodies ineffective.^3-5^ Our results indicate that the mutations in the spike protein of the BA.2.75 sublineage decrease susceptibility to vaccine-induced neutralizing activity compared with BA.2, albeit to a lesser extent than the mutations of BA.4/5. Moreover, BA.2.75 showed an overall higher sensitivity to advanced SARS-CoV-2 neutralizing monoclonal antibodies, including antibodies currently in clinical use. Our results suggest that antibody escape is a less pronounced characteristic of BA.2.75 in comparison to BA.4/5, and additional features favoring viral expansion might be required for BA.2.75 to gain a growth advantage over BA.4/5.

## Figure legends

**Figure S1.**
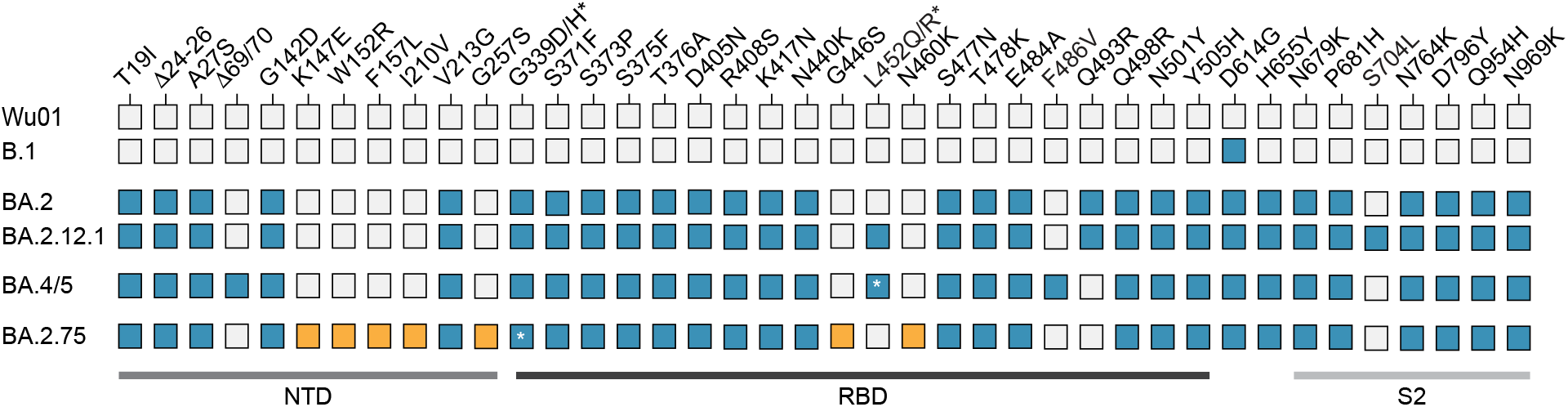
Spike protein mutations. Mutations in the spike protein of SARS-CoV-2 variants relative to the ancestral Wu01 strain are highlighted by colored boxes corresponding to the amino acid changes indicate above. Amongst the tested Omicron sublineages, residues exclusively mutated in BA.2.75 are highlighted in orange. NTD, N-terminal domain; RBD, receptor-binding domain.

**Figure S2.**
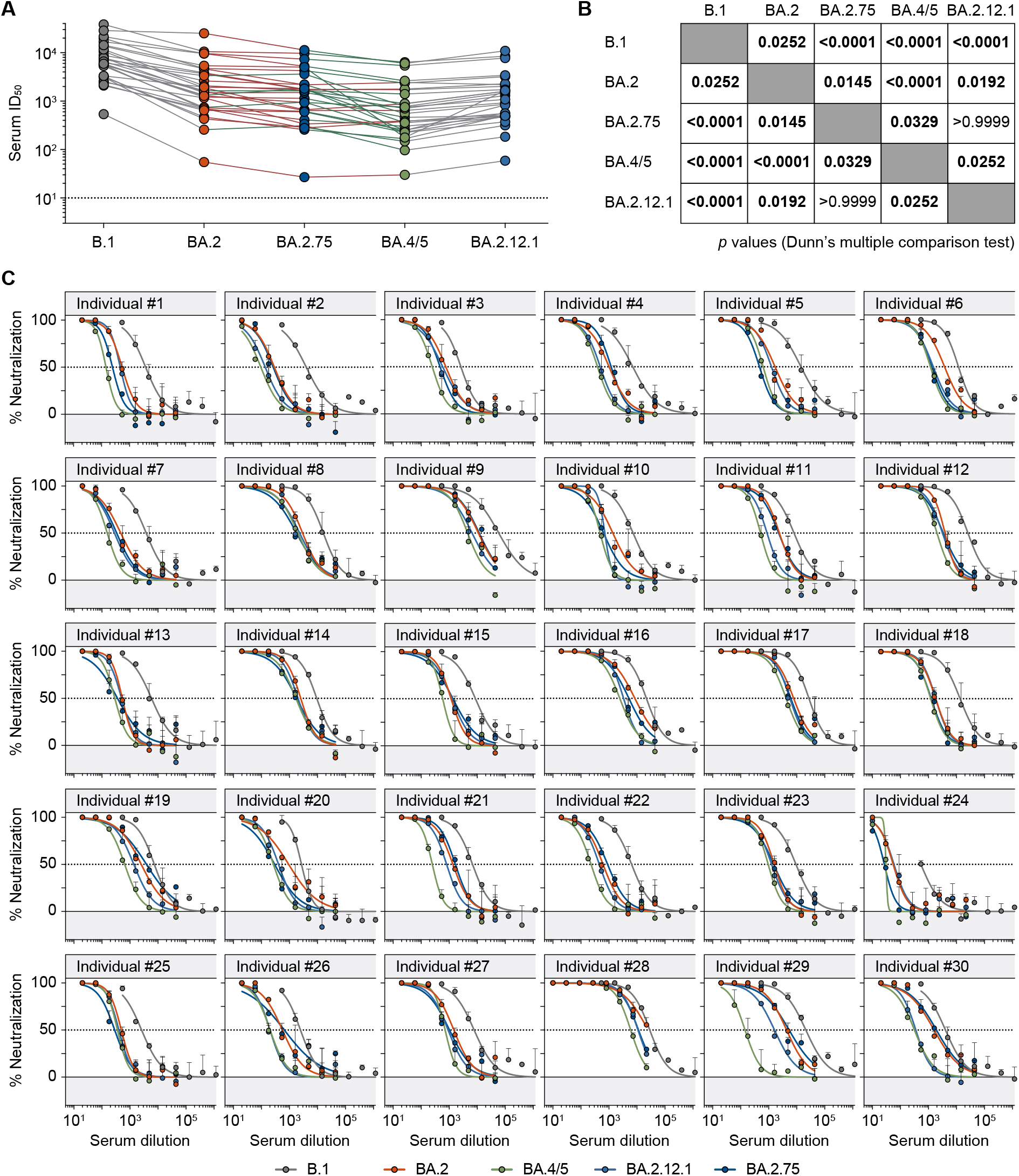
Vaccinee serum neutralizing activity. **(A)** Fifty-percent inhibitory dilutions (ID50s) against SARS-CoV-2 variants in samples collected 4 weeks after a BNT162b2 booster vaccination determined by pseudovirus neutralization assay. Lines connect average serum ID50s (from two experiments for each variant) of individual study participants against the indicated SARS-CoV-2 variants. Lines connecting BA.2 and BA.4/5 to BA.2.75 are color-coded to indicate the ID50 differences relative to BA.2.75 (red lines: BA.2.75 more resistant; green lines: BA.2.75 less resistant). The dotted line indicates the lower limit of quantification (ID50=10). **(B)** Adjusted *p* values of serum ID50 comparisons between indicated variants. Values were calculated using a Friedman test followed by Dunn’s multiple comparison test. Statistically significant differences (adjusted *p* <0.05) are indicated in bold. **(C)** Representative fitted serum SARS-CoV-2 pseudovirus neutralization curves of individual study participants. Circles and bars indicate average values and standard deviation of one experiment. Dotted lines indicate 50% neutralization.

**Figure S3.**
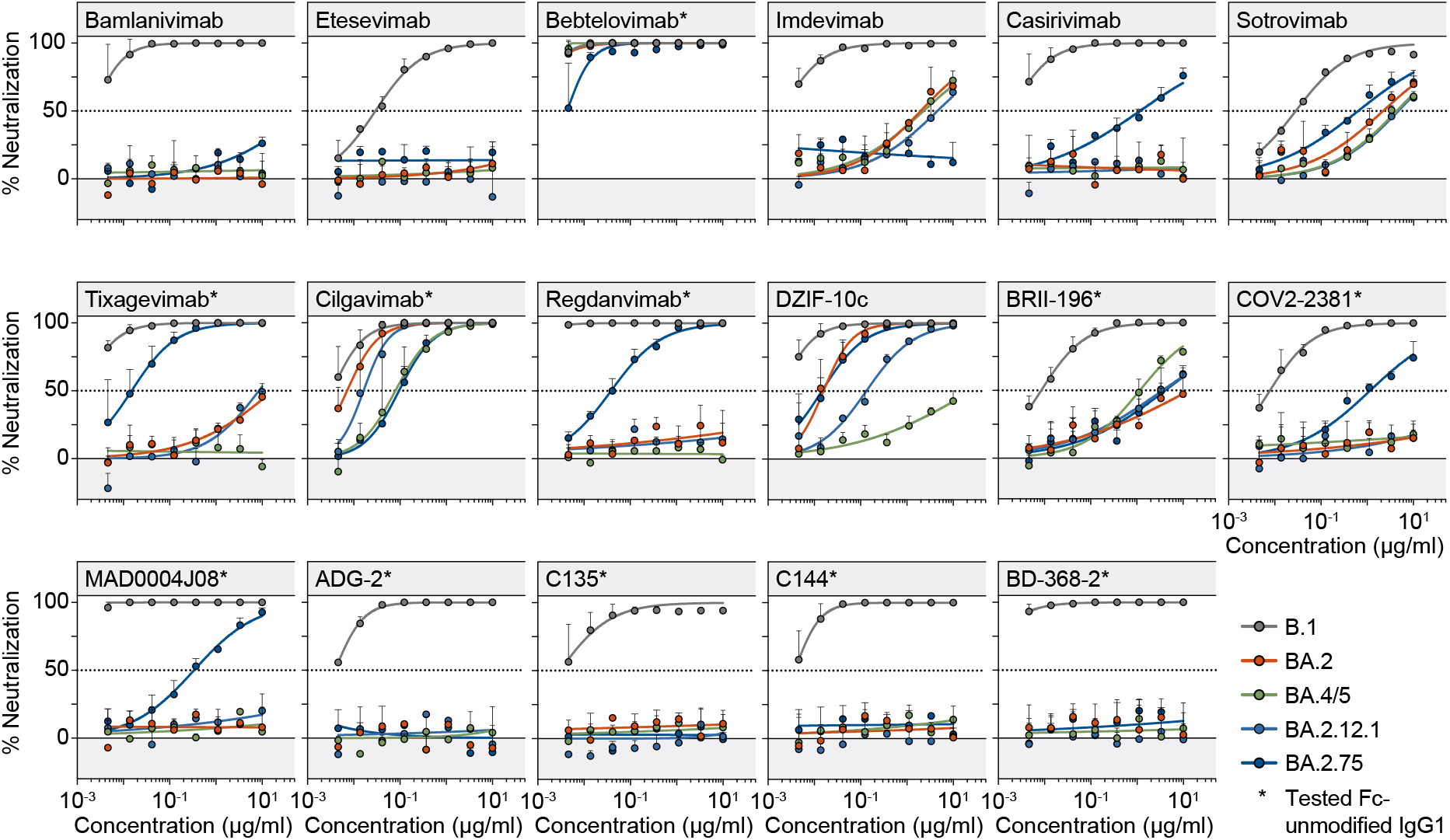
Monoclonal antibody neutralizing activity. Fitted SARS-CoV-2 pseudovirus neutralization curves of monoclonal antibodies. Circles and bars represent average values from two experiments (each performed with technical duplicates) and standard deviation. Dotted lines indicate 50% neutralization.

## Methods

### Study participants and sample collection

Serum samples were collected in two longitudinal cohorts of health care workers and elderly individuals offered the BNT162b2 vaccine as part of routine care ^6,7^. Participants were enrolled at Charité – Universitätsmedizin Berlin, Germany, and at a general practitioner in Berlin, Germany. The study protocol was approved by the ethics committee (EC) of the Federal State of Berlin as well as the Paul Ehrlich Institute (COVIM, EudraCT No. 2021-001512-28). All study participants provided written informed consent. All serum samples were tested to confirm the absence of antibodies targeting the SARS-CoV-2 nucleocapsid protein (SeraSpot Anti-SARS-CoV-2-IgG microarray-based immunoassay, Seramun Diagnostica) that would indicate prior SARS-CoV-2 infection, and no participants with a history of SARS-CoV-2 infection or a positive SARS-CoV-2 nucleic acid amplification test performed at the time of blood collection were included into the analysis. Serum was collected after centrifugation and samples were stored at -80°C until thawing, after which they were stored at 4°C.

### Monoclonal antibodies

Clinical stocks diluted in Dulbecco’s Phosphate Buffered Saline (DPBS, Thermo Fisher) were used to analyze the activity of bamlanivimab, casirivimab, DZIF-10c, etesevimab, imdevimab, and sotrovimab. All remaining monoclonal antibodies were produced as IgG1 isotype antibodies without Fc domain modifications after cloning of codon-optimized antibody heavy and light chain gene segments into expression plasmids (IgG1 and Ig*λ*/Ig*κ*) and co-transfection of heavy and light chain expression plasmids into 293-6E cells (National Research Council of Canada) using 25 kDa branched polyethylenimine (Sigma-Aldrich). After 6-7 days at 37°C and 6% CO_2_ under continuous shaking in FreeStyle Expression Medium supplemented with 20 U/ml penicillin and 20 µg/ml streptomycin (all Thermo Fisher), culture supernatants were harvested, clarified, and incubated with Protein G Sepharose 4 FastFlow (Cytiva) under constant rotation at 4°C overnight. Antibodies were eluted from Protein G beads in chromatography columns (Bio-Rad) after centrifugation and washing with PBS using 0.1 M glycine (pH=3.0) and immediately buffered in 1 M Tris (pH=8.0). Subsequent buffer exchange to DPBS was performed using centrifugal filter units. Antibody heavy and light chain sequences were derived from the following sources: Bebtelovimab (LY-CoV1404) from ref. ^8^; cilgavimab (COV2-2130), tixagevimab (COV2-2130), and COV2-2381 from refs. ^9^ and ^10^; regdanvimab (CT-P59) from ref. ^11^; BRII-196 (P2C-1F11) from ref. ^12^; C135 and C144 from ref. ^13^; MAD0004J08 from ref. ^14^; ADG-2 from ref. ^15^; and BD-368-2 from ref. ^16^.

### Pseudovirus cloning and production

Codon-optimized SARS-CoV-2 spike protein expression plasmids were generated by cloning of gene fragments (Thermo Fisher) and/or site-directed mutagenesis (Q5 Site-Directed Mutagenesis Kit, New England Biolabs). Compared with the Wu01 strain spike protein amino acid sequence, the following changes were included in the respective expression plasmids: B.1: D614G. BA.1: A67V, Δ69-70, T95I, G142D, Δ143-145, N211I, Δ212, ins215EPE, G339D, S371L, S373P, S375F, K417N, N440K, G446S, S477N, T478K, E484A, Q493R, G496S, Q498R, N501Y, Y505H, T547K, D614G, H655Y, N679K, P681H, N764K, D796Y, N856K, Q954H, N969K, and L981F. BA.1.1: As for BA.1 with an additional R346K mutation. BA.2: T19I, Δ24-26, A27S, A67V, G142D, V213G, G339D, S371F, S373P, S375F, T376A, D405N, R408S, K417N, N440K, S477N, T478K, E484A, Q493R, Q498R, N501Y, Y505H, D614G, H655Y, N679K, P681H, N764K, D796Y, Q954H, N969K. BA.2.12.1: As for BA.2 with additional L452Q and S704L mutations. BA.2.75: As for BA.2 with additional K147E, W152R, F157L, I210V, G257S, G339H (D339H relative to BA.2), G446S, and N460K mutations, but lacking the Q493R mutation. BA.4/5: As for BA.2 with additional Δ69-70, L452R, and F486V mutations, but lacking the Q493R mutation. All plasmid spike sequences were verified by Sanger sequencing.

Production of SARS-CoV-2 pseudovirus was performed as previously described.^17,18^ In brief, individual expression plasmids encoding for the SARS-CoV-2 spike protein, HIV-1 Gag/Pol, HIV-1 Tat, HIV-1 Rev, and luciferase were co-transfected in HEK293T cells using the FuGENE 6 Transfection Reagent (Promega) in Dulbecco’s Modified Eagle Medium (DMEM, Thermo Fisher) supplemented with 10% fetal bovine serum (FBS, Sigma-Aldrich), 1 mM sodium pyruvate, 2 mM L-glutamine, 100 IU/ml penicillin, and 100 µg/ml streptomycin (Thermo Fisher). Following a medium exchange one day post-transfection, pseudovirus supernatants were collected between 48 h and 72 h post-transfection, clarified by centrifugation and filtration using a 0.45 µm filter, and stored at -80°C.

### Pseudovirus neutralization assay

The pseudovirus neutralization assay was performed as previously described:^17^ For pseudovirus titration, 293T-ACE2 cells^18^ were co-incubated with serial dilutions of pseudovirus supernatants for two days at 37°C and 5% CO_2_. After the addition of luciferin/lysis buffer (10 mM MgCl_2_, 0.3 mM ATP, 0.5 mM coenzyme A, 17 mM IGEPAL CA-630 (all Sigma-Aldrich), and 1 mM D-Luciferin (GoldBio) in Tris-HCL), relative light units (RLUs) were determined using a microplate reader (Berthold).

All serum samples were heat-inactivated for 45 min at 56°C prior to use. Serial dilutions (1:3) of serum samples (starting at 1:10 or 1:20 for Omicron sublineages and at 1:540 for B.1) or monoclonal antibodies (starting at 10 µg/ml) were co-incubated with pseudovirus supernatants at 37°C and 5% CO_2_ for one hour prior to addition to 293T-ACE2 cells. Following a two-day incubation at 37°C and 5% CO_2_, RLUs were determined as described above. All serum and antibody samples were tested in two experiments each with technical duplicates (with the exception of two serum samples with limited material, which were tested in one experiment with technical duplicates). For the summary of data presented in Figure 1, the average IC_50_ and average ID_50_ are provided.

To determine serum fifty-percent inhibitory dilutions (ID_50_s) and monoclonal antibody fifty-percent inhibitory concentrations (IC_50_s), average background RLUs from uninfected control wells were subtracted and ID_50_s/IC_50_s were determined as the serum dilutions/antibody concentrations resulting in a 50% signal reduction compared to untreated infection control wells. To this end, a non-linear fit model plotting an agonist vs. normalized dose-response curve with variable slope based on the least squares fitting method was used in Prism 7.0 (GraphPad). For one serum sample with a BA.2 ID_50_ above the upper limit of quantification (ID_50_ >21,870), the value was extrapolated based on the neutralization curve.

### Statistical analysis

For the statistical analysis of serum ID_50_s, serum ID_50_s were compared using a Friedman test followed by the Dunn’s multiple comparison test in Prism 7.0 (GraphPad). Differences with adjusted *p* values <0.05 were considered statistically significant.

## Author contributions

Conceptualization, H.G., K.V., and F. Klein; methodology, H.G., K.V., and F. Klein; investigation, H.G. and K.V.; resources and clinical study, P.T-L., D.H., L.E.S., and F. Kurth; formal analysis, H.G. and K.V.; writing-original draft, H.G.; writing-review and editing, K.V., P.T-L., D.H., L.E.S., and F. Klein; visualization, H.G.; supervision, L.E.S., F. Kurth, and F. Klein; funding acquisition, L.E.S., F. Kurth, and F. Klein.

## Acknowledgments

We are grateful to all study participants for their dedication to our research. We thank the members of the COVIM Study Group (Y. Ahlgrimm, L. Bardtke, K. Behn, N. Bethke, H. Bias, D. Briesemeister, C. Conrad, V. M. Corman, C. Dang-Heine, S. Dieckmann, C. Eroglu, D. Frey, J.-A. Gabelich, J. Gerdes, U. Gläser, A. Hetey, L. Hasler, E. T. Helbig, W. Hirst, A. Horn, C. Hülso, S. Jentzsch, C. von Kalle, L. Kegel, A. Krannich, W. M. Koch, P. Kopankiewicz, P. Kroneberg, I. Landgraf, H. Le, L. J. Lippert, C. Lüttke, P. de Macedo Gomes, B. Maeß, C. Matthaiou, J. Michel, F. Münn, A. Nitsche, A.-M. Ollech, C. Peiser, A. Pioch, C. Pley, K. Pohl, C. Rubisch, L. Ruby, A. Sanchez Rezza, I. Schellenberger, V. Schenkel, J. Schlesinger, S. Schmidt, G. Schwanitz, T. Schwarz, J. Seybold, A. Solarek, A. Stege, S. Steinbrecher, P. Stubbemann, C. Thibeault, and D. Treue) for sample acquisition and processing; M. Korenkov for help with antibody cloning; and T. Lange and J.J. Malin for providing clinical antibody stocks. This work was supported by grants from COVIM: NaFoUniMedCovid19 (FKZ: 01KX2021) (to L.E.S. and F. Klein), the Federal Institute for Drugs and Medical Devices (V-2021.3 / 1503_68403 / 2021–2022) (to L.E.S. and F. Kurth), the German Center for Infection Research (DZIF) (to F. Klein), and the Deutsche Forschungsgemeinschaft (DFG) CRC1310 (to F. Klein). The funders had no role in study design, data collection and analysis, manuscript preparation, or the decision to submit for publication.

---

**Table S1.** Study cohort

|  |  |
| --- | --- |
| **Participants - *n*** | 30 |
| **Demographics** |  |
| **Age** |  |
| Median years (IQR; range) | 52 (32-74; 27-96) |
| **Gender** |  |
| Female - *n* (%) | 18 (60%) |
| Male - *n* (%) | 12 (40%) |
| **Reported comorbidities** |  |
| Cardiovascular disease - *n* (%) | 9 (30%) |
| Respiratory disease - *n* (%) | 3 (10%) |
| Immunodeficiencies - *n* (%) | 3 (10%) |
| Diabetes - *n* (%) | 1 (3%) |
| **Vaccination** |  |
| **Vaccine administered** | BNT162b2 |
| **Time between first and second dose** |  |
| Median days (IQR; range) | 21 (21; 21) |
| **Time between second and third dose** |  |
| Median days (IQR; range) | 243 (209-265; 186-286) |
| **Sampling time point** |  |
| **Time of visit after third dose** |  |
| Median days (IQR; range) | 25 (22-29; 14-83) |

